# Homomorphic ZW chromosomes 1 in a wild strawberry show distinctive recombination heterogeneity but a small sex-determining region

**DOI:** 10.1101/029611

**Authors:** Jacob A Tennessen, Rajanikanth Govindarajulu, Aaron Liston, Tia-Lynn Ashman

## Abstract

- Recombination in ancient, heteromorphic sex chromosomes is typically suppressed at the sex-determining region (SDR) and proportionally elevated in the pseudoautosomal region (PAR). However, little is known about recombination dynamics of young, homomorphic plant sex chromosomes.
- We examine male and female function in crosses and unrelated samples of the dioecious octoploid strawberry *Fragaria chiloensis* in order to map the small and recently evolved SDR controlling both traits and to examine recombination patterns on the incipient ZW chromosome.
- The SDR of this ZW system is located within a 280kb window, in which the maternal recombination rate is lower than the paternal. In contrast to the SDR, the maternal PAR recombination rate is much higher than the rates of the paternal PAR or autosomes, culminating in an elevated chromosome-wide rate. W-specific divergence is elevated within the SDR and a single polymorphism is observed in high species-wide linkage disequilibrium with sex.
- Selection for recombination suppression within the small SDR may be weak, but fluctuating sex ratios could favor elevated recombination in the PAR to remove deleterious mutations on the W. The recombination dynamics of this nascent sex chromosome with a modestly diverged SDR may be typical of other dioecious plants.

## Introduction

Sex chromosomes display distinctive inheritance patterns, effective population sizes, recombination frequencies, and structural arrangements, giving them a key role in numerous biological phenomena (Bull, 1983; Charlesworth & Mank, 2010). Sex chromosome systems include male heterogamety (XY inheritance), female heterogamety (ZW inheritance), and other systems (Bachtrog *et al*., 2014). Although sex phenotype is typically thought to be determined by only one or two genes (Bachtrog *et al*., 2014), these functional element(s) frequently reside within a larger sex-determining region (SDR), inherited as a unit. The remainder of the sex chromosome outside the SDR is known as the pseudoautosomal region (PAR) (Ohno, 1969; Otto *et al*., 2011). A defining feature of sex chromosomes is a specific and repeatable pattern of heterogeneity in recombination rate, seen across diverse taxa (Bergero & Charlesworth, 2009). The heterogametic parent typically shows suppressed recombination in the SDR and elevated recombination in the PAR (reviewed in Otto *et al*., 2011; Charlesworth, 2015). The adaptive basis for these recombination patterns has been extensively examined theoretically (Charlesworth & Charlesworth, 1978; Bachtrog *et al*., 2014). In brief, if an SDR-linked locus harbors an allele that is advantageous in one sex and disadvantageous in the other (sexually antagonistic), recombinants will have decreased fitness and recombination suppression will be favored. In the case where sex is determined by two loci both carrying fertility/sterility mutations, recombinants lack both fertility alleles and are thus neuters, making the suppression of recombination highly advantageous (Bull, 1983; Charlesworth & Charlesworth, 1978; Spigler *et al*., 2008). As recombination is suppressed, differential substitutions and insertions accumulate within the widening SDR, resulting in evolutionary strata of sequence divergence between X/Z and Y/W (Bergero & Charlesworth, 2009). The strength of selection for expansion and divergence of the SDR likely depends on the degree of sexual dimorphism, and therefore the number of genes with alleles showing sex-specific fitness (Ahmed *et al*., 2014). Recombination is nonetheless usually required for faithful chromosome segregation, producing an increased PAR recombination rate in the heterogametic sex because equivalent recombination occurs over a smaller physical distance (Otto *et al*., 2011). Consistent with this hypothesis, the total sex chromosome recombination rate averaged across the PAR and SDR is usually less than or equal to the genome-wide average (Otto *et al*., 2011). These characteristic recombination dynamics of sex chromosomes are the evolutionary basis for their other unusual features such as heteromorphy, rapid molecular evolution, degeneration, and determination of hybrid incompatibility (Charlesworth, 1996).

The most extensively studied sex chromosomes are ancient and highly heteromorphic, and thus provide little insight into the initial stages of sex chromosome evolution (Bernardo *et al*., 2009; Cortez *et al*., 2014). However, a much greater diversity of sex determining systems exists, especially in plants (Diggle *et al*., 2011; Bachtrog *et al*., 2014; Charlesworth, 2015). While few dioecious flowering plants have heteromorphic sex chromosomes or even known sex-determination mechanisms (Charlesworth & Mank, 2010; Ming *et al*., 2011; Renner, 2014), several recent studies have revealed homomorphic sex chromosomes with small SDRs, including in *Vitis* (143kb; Fechter *et al*., 2012; Picq *et al*., 2014) and *Populus* (100kb; Geraldes *et al*., 2015). The slightly larger SDRs (1-10Mb) in plant taxa such as *Actinidia* (Zhang *et al*., 2015), *Carica* (Wang *et al*., 2012; Lappin et al. 2015), and *Asparagus* (A. Harkess, personal communication) still encompass small ≤10%) proportions of their chromosomes, and are much smaller than cytogenetically heteromorphic chromosomes. Despite PAR characterizations in older systems (e.g. Qiu et al. 2016; Lappin et al. 2015), no study has comprehensively examined chromosome-wide recombination heterogeneity in a plant with a very small (<500kb) SDR. Furthermore, most previous studies in plants have examined XY systems in diploids (but see Pucholt *et al*., 2015; Russell & Pannell, 2015), although ZW systems may have distinct dynamics (Ming *et al*., 2011), and the potential association between dioecy and polyploidy (Miller & Venable, 2000; Ashman *et al*., 2013) makes polyploids an attractive study system. Young sex chromosomes are ideal models for studying evolutionary processes related to sex (Bergero *et al*., 2013), and indeed could represent the evolutionary precursors of heteromorphic chromosomes. Alternatively, SDRs may remain relatively small for tens of millions of years (Zhou *et al*., 2014), or control of sex may turn over so rapidly among distinct genomic regions that SDRs undergo minimal evolutionary change (Dufresnes *et al*., 2015). Recombination rates are determined by several factors that are only partially understood (Bomblies *et al*., 2015; Shilo *et al*., 2015).

Early sex chromosome evolution has progressed particularly rapidly and variably in *Fragaria* (Spigler *et al*., 2008; Ashman *et al*., 2012; Liston *et al*., 2014). Evolving from a hermaphroditic ancestor just 2 mya (Njuguna *et al*., 2014), the strawberry genus has diversified into species displaying sexual systems including gynodioecy (females and hermaphrodites), subdioecy (females, males, and hermaphrodites), and dioecy (females and males). In the gynodioecious diploid *F. vesca* subsp. *bracteata*, at least two unlinked loci harbor male sterility alleles, only one of which, on chromosome IV (out of seven homeologous chromosome groups, designated I-VII; Table 1), is heterozygous in females (Tennessen *et al*., 2013; Ashman *et al*., 2015). *Fragaria vesca* subsp. *bracteata* is an ancestor of allo-octoploid *Fragaria* species (2n=8x=56), having provided both cytoplasmic DNA and the Av subgenome (one of four nuclear subgenomes; Table 1) (Njuguna *et al*., 2014; Tennessen *et al*., 2014; Govindarajulu *et al*., 2015; Sargent *et al*., 2016). It does not share its sex loci with the octoploid clade, though. Octoploid *F. virginiana* subsp. *virginiana* possesses a female-heterozygous SDR at the start of chromosome VI-B2-m (i.e., homeologous group six, subgenome B2, maternal chromosome), while octoploid *F. chiloensis* possesses a female-heterozygous SDR at the end of chromosome VI-Av-m, here designated “ZW” (Spigler *et al*., 2010; Spigler *et al*., 2011; Goldberg *et al*., 2010; Tennessen *et al*., 2014; Table 1). Octoploid *Fragaria* are highly diploidized, such that the subgenomes remain evolutionarily distinct (Tennessen *et al*., 2014). Thus, sex determination in this genus has frequently evolved independently, and/or exhibits a rapidly shifting genomic position. Moreover, since the octoploid clade arose 1 mya (Njuguna *et al*., 2014), and SDRs are not shared among the dimorphic species they appear to be some of the youngest studied to date.

**Table 1.**
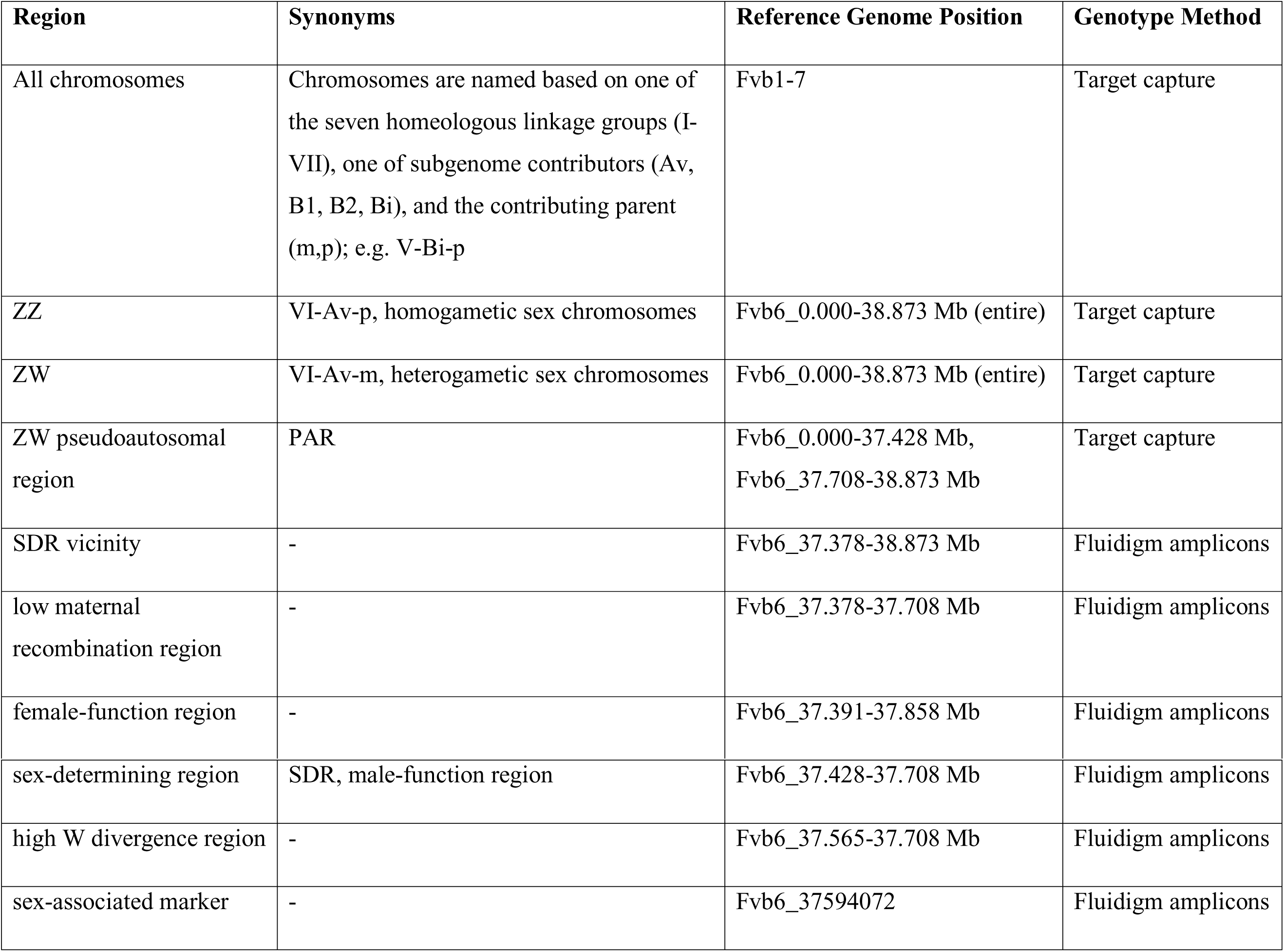
Regions of the *F. chiloensis* genome relevant to sex determination, showing notations used in text, synonyms, locations in reference genome, and methods used to characterize them.

*Fragaria chiloensis* occurs along the Pacific coasts of North and temperate South America. The species is nearly completely dioecious, although some hermaphrodites do occur (Hancock & Bringhurst, 1980), and has undergone perhaps the greatest sexual differentiation in the genus with respect to the hermaphroditic *Fragaria* ancestor (Ashman, 2005; Ashman *et al*., 2012). Thus, although sex phenotypes (Spigler & Ashman 2011; Spigler *et al*., 2011) and genomics (Shulaev *et al*., 2011; Sargent *et al*., 2016) of other *Fragaria* species have been studied more extensively, F. *chiloensis* is a more complete sex chromosome model. Because male and female function both map to a single maternal genomic location, with no previously observed recombination between them (Goldberg *et al*., 2010), *F. chiloensis* has all the hallmarks of a complete, albeit incipient, ZW sex chromosome. It is possibly the youngest ZW system known in plants, similar in age to the youngest XY systems. As an octoploid, it is ideally suited for recombination analysis because recombination rates can be compared among subgenomes and between the sex chromosome and autosomal homeologs. However, neither the precise genomic location of the SDR, its physical size, nor the recombination profile of the SDR-carrying chromosomes could be determined from the initial microsatellite-based linkage map (Goldberg *et al*., 2010). Here we characterize the *F. chiloensis* sex chromosome with respect to Z-or W-specific recombination rates in the PAR and SDR, as well as recombination-dependent metrics including sequence divergence and linkage disequilibrium, in order to infer the initial evolutionary adjustments in recombination rate and divergence that occur in response to sex linkage.

## Materials and Methods

### Samples

Our primary cross (HM1×SAL3) represents an expanded set of F1 offspring from maternal *Fragaria chiloensis* subsp. *lucida* (HM1, previously GP33) and paternal *Fragaria chiloensis* subsp. *pacifica* (SAL3) used previously (as GP×SAL, Goldberg *et al*., 2010; Table 2). Female HM1 was obtained from the USDA National Clonal Germplasm Repository (Accession PI 612489; originally collected from Honeyman Memorial State Park, Oregon, 43.93N, 124.11W). Hermaphrodite (10% female fertility) SAL3 from Salishan, Oregon (44.91N, 124.02W) was chosen to assess paternal inheritance of female fertility. We hand-pollinated pistillate HM1 flowers with SAL3 pollen. We planted 1695 seeds in sets across five years (between 2009 and 2015). We transplanted *c*. 2-month-old seedlings into 8-cm-square pots with a 2:1 mixture of Fafard #4 (Conrad Fafard) and sand. After germination the plants initially received 513 mg granular Nutricote 13:13:13 N:P:K fertilizer (Chisso-Asahi Fertilizer). Plants were grown under 15°:20° C night:day temperatures and 10 to 12 h days throughout the majority of the flowering period. To initiate flowering plants were exposed to 8°: 12° C night:day temperatures with an 8 h low light day. Additional liquid fertilizer and pest control measures were applied as needed.

**Table 2.** Details of all three *F. chiloensis* crosses, including relative divergence of Z and W haplotypes in the SDR vicinity.

| **Cross** | **Maternal Parent** | **Paternal Parent** | **Offspring number** | **SNPs in coupling, SDR vicinity<sup>a</sup>** | **SNPs in coupling, high W divergence window<sup>b</sup>** | **SNPs in repulsion, SDR vicinity<sup>c</sup>** | **Paternally segregating, SDR vicinity<sup>d</sup>** |
| --- | --- | --- | --- | --- | --- | --- | --- |
| HM1×SA L3 | HM1 (aka GP33<sup>e</sup>) | SAL3 | 1275 | 18 | 10 (2 nonsynonymous, 1 synonymous, 7 noncoding<sup>f</sup>) | 18 | 9, 15 |
| EUR13× EUR3 | EUR13 | EUR3 | 20 | 2 | 1 (1 noncoding<sup>f</sup>) | 2 | 0, 3 |
| PTR14×P TR19 | PTR14 | PTR19 | 20 | 7 | 4 (2 nonsynonymous, 2 noncoding<sup>f</sup>) | 5 | 0, 4 |

<sup>a</sup>Number of maternal SNPs in coupling male sterility/female fertility (i.e. on W) in the SDR vicinity807 <sup>b</sup>Number of maternal SNPs in coupling male sterility/female fertility within the high-W divergence window
<sup>c</sup>Number of maternal SNPs in repulsion male sterility/female fertility (i.e. on Z) in the SDR vicinity
<sup>d</sup>Number of paternal SNPs segregating on each of two haplotypes (i.e. on either Z) in the SDR vicinity
<sup>e</sup>See Material and Methods and Goldberg *et al.* (2010)
<sup>f</sup>One noncoding “sex-associated marker” is shared among all three crosses

We also examined the parents and F1 offspring of two other independent *F. chiloensis* crosses, in order to test the universality of the SDR location identified in HM1×SAL3 and to identify single nucleotide polymorphisms (SNPs) universally on the SDR W haplotype (Table 2). The parents of the first cross (EUR13×EUR3) were a female (EUR13) and a male (EUR3; 0% female fertility) *F. chiloensis* subsp. *lucida* from Eureka, California (40.762 N, 124.225 W). The parents of the second cross (PTR14×PTR19) were a female (PTR14) and an hermaphrodite (PTR19; 10% female fertility) *F. chiloensis* subsp. *lucida* from Point Reyes, California (38.0683 N, 122.971 W). We raised 105 and 65 F1 offspring from the two crosses, respectively, following the same planting and growth regimes as for HM1×SAL3. As the goal of these crosses was validation of the sex-linked SNP results from HM1×SAL3, rather than fine-mapping, we examined substantially fewer offspring (20 each) than for HM1×SAL3.

We also examined 16 unrelated plants from six populations (including the four source populations of cross plants) across the geographic range of *F. chiloensis* (Table S1). The plants were collected from the wild as clones or obtained from the National Clonal Germplasm Repository and raised under the same planting and growth conditions.

### Sex phenotyping

In order to define the SDR, we first phenotyped male and female function following Spigler *et al*., (2011). We scored male function, on at least two flowers per plant two separate times, as a binary trait: “male fertile” if possessing large, bright-yellow anthers that visibly released pollen, and “male sterile” if possessing vestigial white or small, pale-yellow anthers that neither dehisced nor showed mature pollen. Ambiguous anthers were examined microscopically for mature pollen. To ensure full potential fruit set, we hand pollinated plants three times weekly with outcross pollen. We estimated female fertility as a continuous quantitative trait for each individual as the proportion of flowers that produced fruit (fruits:total pollinated flowers) following Spigler *et al*., (2011). We calculated the association between male function and female function using simple linear regression (R Core Team, 2013). Because female fertility may be influenced by plant size (estimated as total number of flowers or buds produced), we also included total flower number as a second independent variable in multiple regression (R Core Team, 2013). For HM1×SAL3 female fertility was treated as continuous, while for the EUR13×EUR3 and PTR14×PTR19 crosses and the unrelated individuals, it was converted to a binary trait following Goldberg *et al*., (2010).

### DNA extraction and quantification

Fresh tissue DNA extractions of 42 HM1×SAL3 offspring were described previously (Tennessen *et al*., 2014). For the remaining plants, we extracted DNA from leaf tissue stored on silica gel, using the Norgen biotek plant/fungi high-throughput 96 well DNA isolation kit (Norgen Biotek, ON, Canada) and the DNA extraction service provider Agbiotech Inc (CA, USA). We added an additional 100ul 10% SDS and 10ul β-mercaptoethanol to the lysis buffer to improve DNA yield. DNA was eluted with sodium acetate and ethanol precipitated. We performed Picogreen assays using a Tecan plate reader and diluted DNA to 50ng:ul using DEPC treated water for microfluidics PCR on the Fluidigm access array system (https://www.fluidigm.com/products/access-array).

### Target capture mapping

We previously described a target capture linkage map of 2542 segregating *F. chiloensis* markers generated using 42 HM1×SAL3 offspring (Tennessen *et al*., 2014). All markers align to a distinct position in the *F. vesca* reference genome (v. 2.0, here designated “Fvb” as in Tennessen *et al*., 2014), and have a linkage map position (cM) on a paternal (“p”) or maternal (“m”) linkage group pertaining to one of the four subgenomes (Table 1). In order to map sex phenotypes, we first added binary male function to the linkage map with OneMap (Margarido *et al*., 2007), as in Tennessen *et al*., (2013). Because female fertility is a continuous trait (Fig 1), we identified female function quantitative trait loci (QTL) using R/qtl (Broman *et al*., 2003; R Core Team, 2013). We used the scanone option to identify possible female function QTL (minimum LOD 3). We checked whether a multiple additive QTL model was supported, with a LOD at least 3 greater than the LOD of the single best QTL, using the scantwo option. The Fvb region where both male function and female function QTL mapped, presumably harboring the SDR and possibly also adjacent portions of the PAR, was designated the “SDR vicinity” and examined with additional genotyping (Table 1).

**Fig 1.**
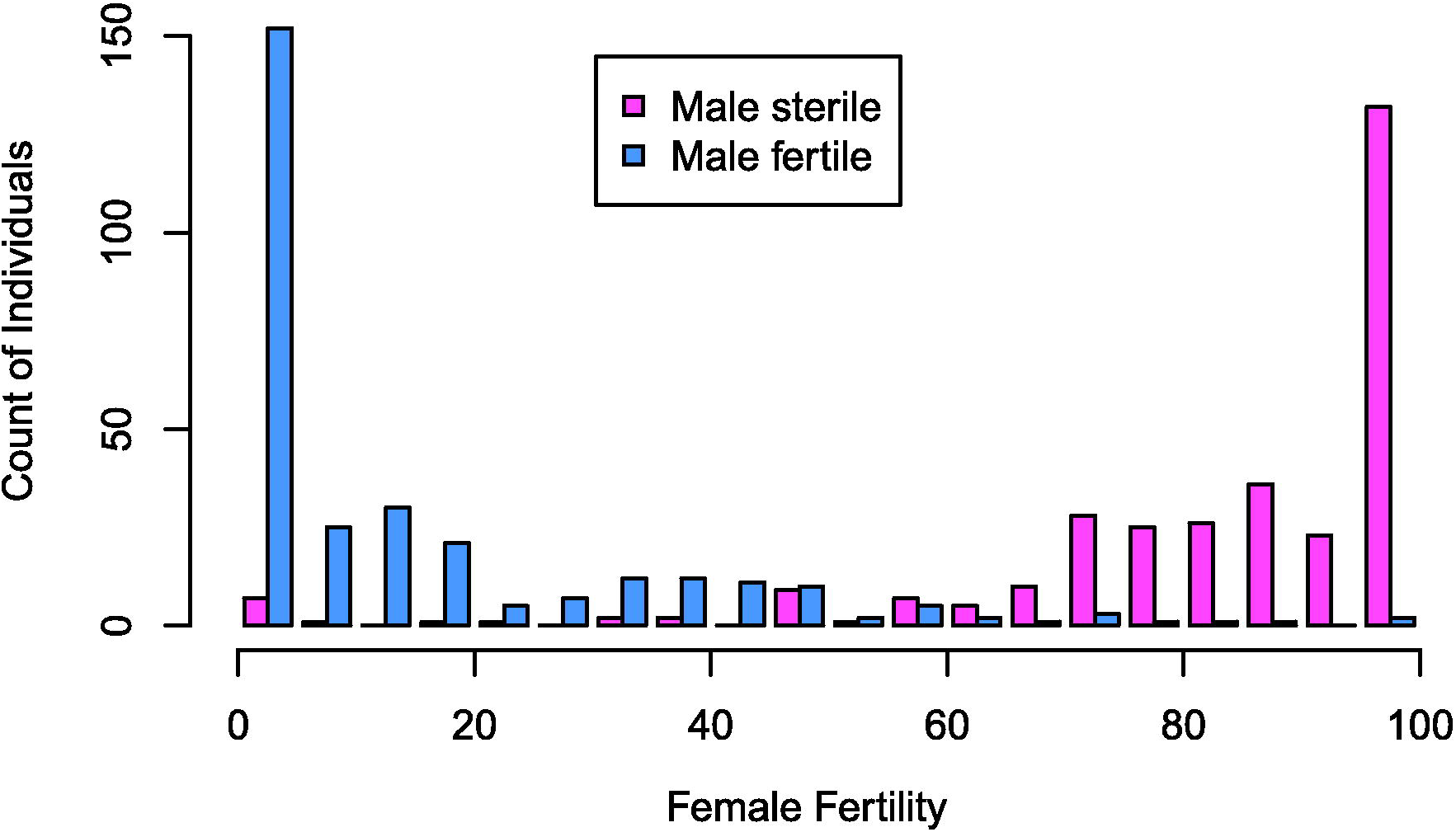
Sex phenotype distributions among *Fragaria chiloensis* offspring of HM1×SAL3 cross. Male function is defined as the ability (male fertility, blue) or inability (male sterility, pink) to produce mature pollen and is a binary trait found in approximately half of the progeny. Female fertility is defined as the proportion of flowers producing fruit, and has a continuous though bimodal distribution, shown here partitioned into bins of 5%. Male and female function are highly negatively correlated, with nearly all male-fertile offspring showing <50% female fertility, and nearly all male-sterile offspring showing >50% female fertility.

In order to estimate genome-wide recombination rates, we re-examined the target capture HM1×SAL3 linkage maps (Tennessen *et al*., 2014), excluding putative recombination events that could be caused by genotyping error, as in Tennessen *et al*., (2013). We also identified all inconsistencies in marker order relative to Fvb supported by more than one genotype, and considered these to be potential genomic rearrangements. We assumed that physical distances on the octoploid chromosomes could be approximated by Mb distances in Fvb, and thus we estimated recombination rates genome-wide and per each linkage group as cM Mb^−1^. We divided all 56 linkage groups into 5-10Mb segments free from rearrangements, and calculated recombination rate for each segment.

### Amplicon generation and sequencing

In order to fine-map the SDR vicinity and examine recombination rates, we genotyped SDR-vicinity SNPs using Illumina sequencing of PCR amplicons (Cronn *et al*., 2012). The Fluidigm microfluidic approach is an effective method for mapping SNP markers in a large sample of polyploids (Byers *et al*., 2012; Uribe-Convers *et al*., 2016). We performed sequencing in three sequential rounds, refining the combination of amplicons each time based on amplicon performance and fine-mapping results (Table S2; Table S3). Some samples were included in multiple rounds due to poor-quality genotypes or in order to further fine map recombinants. Initially, we designed primers for 48 amplicons of *c*. 300-600bp (larger amplicons performed poorly), with a Fvb position within the SDR vicinity. Most amplicons (41) contained at least one SNP (on any subgenome) identified in the target capture map. Following the Fluidigm Access Array System user guide for Illumina (https://www.fluidigm.com), primer pairs were validated using DNA from HM1 with the Agilent bioanalyzer (Agilent Technologies Inc.) to verify the expected product size and ascertain that on-target products account for >50% of the total yield. Via IBEST at the University of Idaho, we used the Fluidigm 48.48 Access Array IFC (Integrated Fluidic Circuits) to generate amplicons in parallel in microfluidic reactors. Forward and reverse primers were attached with common sequence tags (CS1 & CS2) to enable the addition of P5 and P7 Illumina adapters along with dual index multiplex barcodes. We multiplex sequenced 48 pools of amplicons (one per individual) in 10-12 access array plates on one-half lanes of the Illumina MiSeq, yielding 300-bp paired-end reads. For the first round we sequenced 478 HM1×SAL3 offspring, HM1, and SAL3. For the second round, we retained 19 of the original 48 primer pairs and replaced 29 with new pairs. We used these to sequence 469 HM1×SAL3 offspring, HM1, SAL3, and the parents and offspring from EUR13×EUR3 and PTR14×PTR19. For the third round, we retained 39 primer pairs and replaced 9 with new pairs. We used these to sequence 545 HM1×SAL3 offspring, HM1, SAL3, all EUR13×EUR3 and PTR14×PTR19 individuals, and 18 additional unrelated plants (16 yielding useable genotypes). In total, 1302 unique plants were included in amplicon sequencing. Including the target capture samples, 1274 HM1×SAL3 offspring were genotyped.

### Amplicon genotyping and fine mapping

We called genotypes using POLiMAPS (Tennessen *et al*., 2014). We first processed Illumina reads with dbcAmplicons (https://github.com/msettles/dbcAmplicons), trimmed them with Trimmomatic (Bolger et al., 2014), aligned them to the amplicon regions extracted from Fvb with BWA (Li & Durbin, 2009), and generated pileup files with SAMtools (Li *et al*., 2009). We generated pileup files as in the target capture maps (Tennessen et al., 2014), only including SNPs heterozygous in a single parent. Because HM1×SAL3 amplicon sample sizes were larger and there were more missing genotypes than in the target capture maps, we allowed up to 200 missing genotypes per site and required 40 observations each of homozygotes and heterozygotes for each SNP. For EUR13×EUR3, PTR14×PTR19, and the set of unrelated plants, all with much smaller sample sizes, we allowed up to 10 missing genotypes and required 4 observations each of homozygotes and heterozygotes. We converted genotypes to OneMap format and generated linkage maps in OneMap (LOD of 3).

We identified recombination events to estimate recombination rates in the SDR vicinity and to fine map sex QTL. Because male function showed a near-perfect match to genotype, and the few exceptions were not recombinants, we simply defined the male-function region between the closest recombination events upstream and downstream that caused mismatches between genotype and male function. Because female function is a continuous trait showing a weaker correlation with genotype, we calculated LOD scores for all sites in the SDR vicinity, calculated the correlation between genotype and female function across the SDR vicinity, and used R/cocor (R Core Team, 2013; Diedenhofen & Musch, 2015) to compare these correlations. We defined a female-function region in which correlations were not significantly different from that at the site with the best LOD score. Because male function could be more precisely mapped than female function, we considered the male-function region to comprise the maximum potential extent of the SDR. We examined genes within the male-function region by consulting both the original annotation of Fvb (Shulaev *et al*., 2011) and a re-annotation (Darwish *et al*., 2015). As with the target capture maps, we estimated recombination rates as cM Mb^−1^.

We measured W-specific divergence per cross (not species-wide) as the proportion of SNPs in coupling with male sterility/female fertility among all sequenced sites (using autosomal homeologs and Fvb as outgroups). Our study design precludes examining divergence shared by all Z chromosomes but not W, since such Z-specific SNPs will be homozygous on linkage group VI-Av-p (here designated “ZZ”) and thus will not segregate in the offspring.

## Results

### Phenotypes

In HM1×SAL3, 1285 seeds germinated. We obtained genotype and/or phenotype data from 1275 of these (Table S4). We determined male function for 693 progeny (352 male sterile, 341 male fertile), and female function for 619 progeny (157 with female fertility <5% and thus “female sterile” as defined in Goldberg *et al*., 2010; mean flower number = 15.5). Male and female function were negatively correlated (R^2^ = 0.75; *P* < 10^−15^; Fig 1). Including total flower number (proxy for plant size) in multiple regression analysis did not significantly improve the proportion of variance explained in the simple linear regression (R^2^ = 0.76; *P* < 10^−6^). Similar phenotype distributions were observed among two sets of 20 offspring from independent crosses (EUR13xEUR3 and PTR14×PTR19, Table S5), and among 16 additional unrelated plants (Table S1).

### Target capture mapping

In the HM1×SAL3 target capture dataset (Table S6), we observed 23 male-fertile offspring and 19 male-sterile offspring. Male function mapped to the end of ZW, as expected (LOD = 10.6; Fig 2; Goldberg *et al*., 2010). Specifically, the last recombination event before the male-function region occurred after position Fvb6_37.391Mb, after which only a single male-sterile individual (not recombinant anywhere on the linkage group) had a genotype mismatching sex (R^2^ = 0.91; *P* < 10^−15^). Thus, the SDR occurs somewhere within the remaining 1.482Mb of the chromosome (the “SDR vicinity”), on which we observed 10 SNPs in 9 targeted regions. An alternate, much weaker potential QTL on linkage group II-Av-p (LOD = 3.6; Fig 2) was assumed to be a statistical artifact and not pursued further, because it showed no significant association after accounting for the QTL on ZW (R^2^ = 0.04; *P* > 0.1). The highest QTL for female function (R^2^ = 0.77; *P* < 10^−13^), treated as a quantitative trait, occurred at the same location on ZW as male function, peaking at Fvb6_37.391Mb (LOD = 18.8). The best two-QTL model for female function was not significantly better than the single-QTL model.

**Fig 2.**
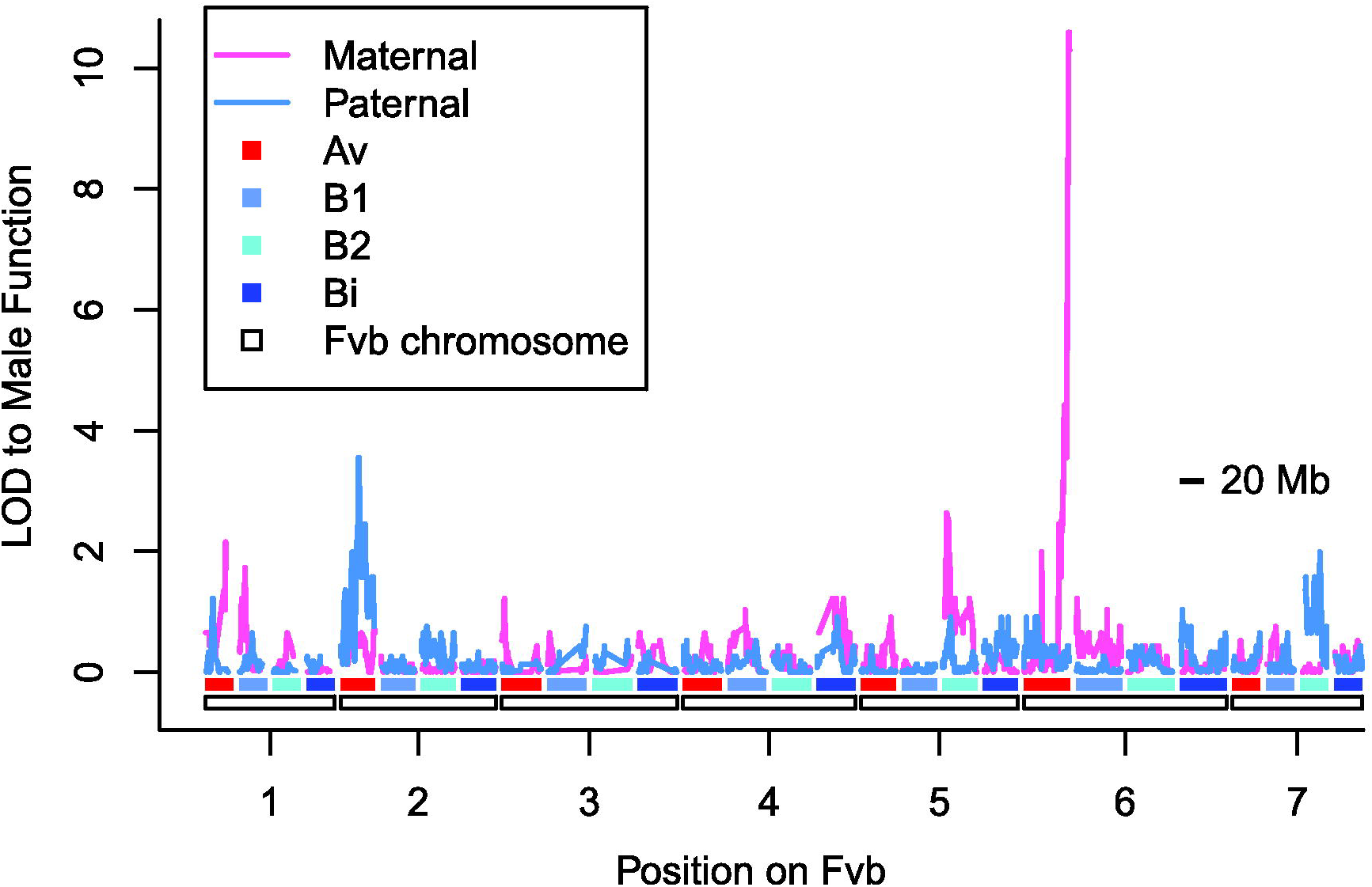
Male function LOD scores across the *Fragaria chiloensis* target capture linkage maps. The haploid chromosomes of the Fvb reference genome are denoted by white rectangles along the x-axis. Each haploid chromosome occurs on each of four subgenomes (Av, B1, B2, or Bi), shown as red or blue shaded bars. LOD scores for maternal and paternal linkage maps are shown on the y-axis. The highest LOD score occurs on the ZW linkage group VI-Av-m, peaking at 10.6 for the last ten segregating sites in the linkage group, all aligning after Fvb6_37.391Mb. We subsequently validated this region on ZW using hundreds of additional offspring, and thus the second-highest LOD peak on linkage group II-Av-p (LOD = 3.6) appears not to be biologically meaningful.

### Amplicon fine mapping

Of the 1285 HM1×SAL3 progeny, we obtained useable SDR-vicinity amplicon genotypes from 1235 offspring, including 1215 Av-m genotypes and 1196 Av-p genotypes (Table S4). Although the set of useable sites varied slightly among each of three sequencing rounds, by the third round we could genotype 36 segregating sites among 24 amplicons on ZW, and 24 segregating sites among 16 amplicons on ZZ. Recombinants were observed in all growing years and all sequencing rounds. We observed 32 plants that recombined on ZW in the SDR vicinity, and these allowed us to further fine-map the male-function region to a 280kb section between Fvb_37.428Mb and Fvb6_37.708Mb (Fig 3; Table 1). Specifically, we observed near-perfect matches to male function at all sites between Fvb6_37.566Mb and Fvb6_37.682Mb, with two male-fertile downstream recombinants showing mismatched genotypes starting at Fvb6_37.708Mb, and two upstream recombinants at Fvb_37.428Mb, one of which was male fertile. Seven individuals had genotypes mismatching male function in this region. These individuals were non-recombinant across the entire SDR vicinity. With <1% of offspring mismatching, the correlation between genotype in this male-function region and male function phenotype was very strong (R^2^ = 0.96; *P* < 10^−15^). The male-function region resides entirely within a single scaffold of Fvb, scf0513124_6, minimizing the possibility of genome assembly errors in this region. The *F. vesca* genome annotations identify 70 genes in this male-function region (Table S7).

**Fig 3.**
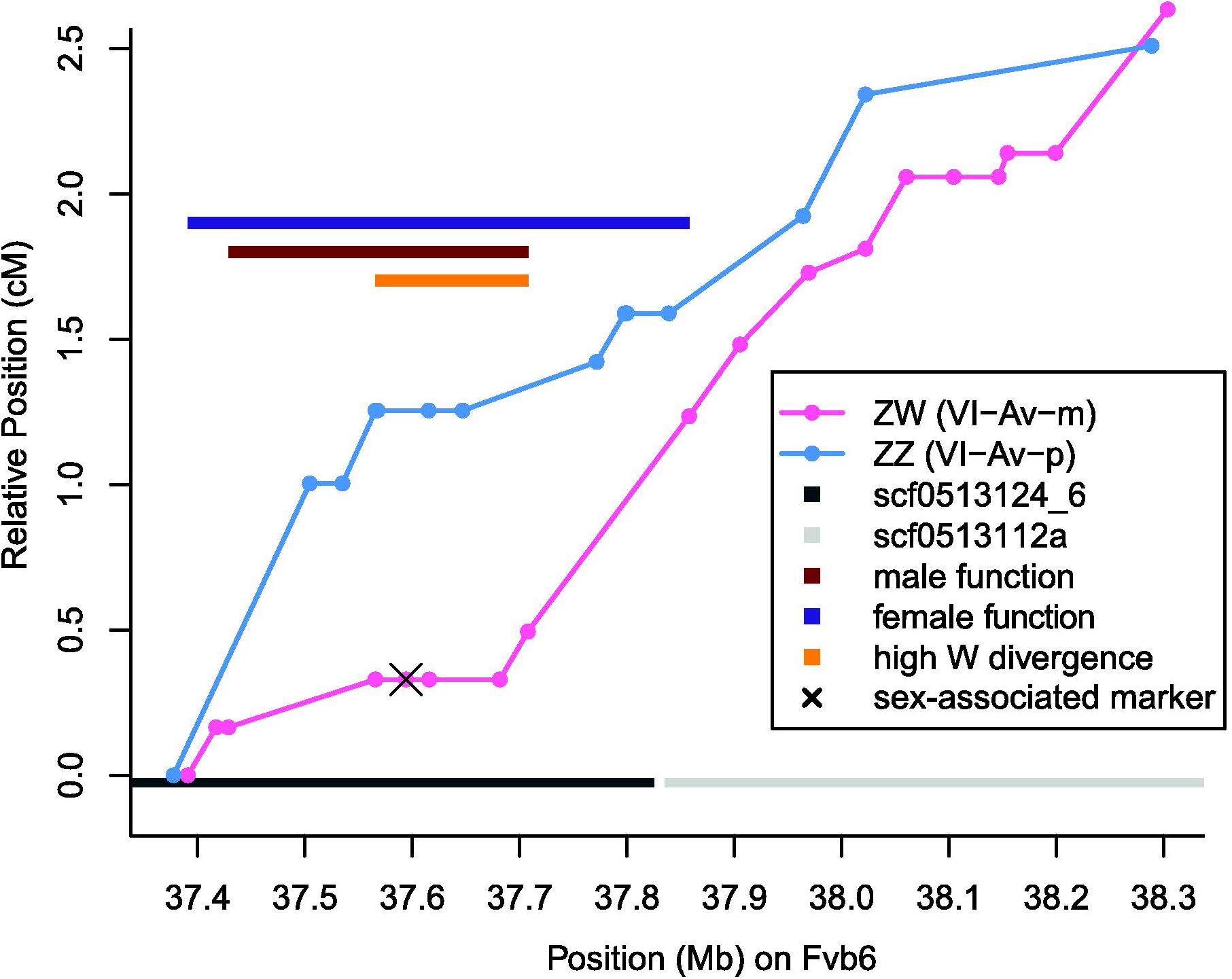
Fine-mapped male and female function and recombination rates across the sex determining region (SDR) vicinity of *Fragaria chiloensis*. Physical distance (Mb) along the distal end of Fvb6, encompassing portions of two genomic scaffolds (scf0513124_6 and scf0513112a), is shown on the x-axis. All amplicon markers in the SDR vicinity in ZW (pink) and ZZ (light blue) are plotted, with relative linkage map position (cM, arbitrarily starting at 0) shown on the y-axis. An X marks the position of the “sex-associated marker” SNP on ZW which shows a high correlation with sex across the full set of unrelated plants. Recombination rates can be inferred from the slopes of lines connecting ZW (pink) and ZZ (light blue) markers. Three colored bars with arbitrary vertical position indicate different estimates of the SDR. The female-function region showing significantly higher correlation with female fertility than the surrounding regions is indicated with a dark purple bar. The male-function region, bounded by the recombinants which decrease the correlation with male function, is indicated with a dark red bar. The high W divergence region, within which most W-specific SNPs are observed, is indicated by an orange bar. The SDR encompasses a relatively small portion of the 39Mb chromosome. Maternal and paternal recombination rates are similar, except near the SDR, where they are lower in the mother.

The highest LOD value for our female function measure (145.9) was observed on ZW at Fvb6_37.428Mb, and correlations that were not significantly weaker than this were observed between Fvb6_37.428-37.708Mb (LOD = 143.1-145.9), coinciding precisely with the male function-region. Significantly worse correlations were observed upstream at Fvb6_37.391Mb (LOD = 142.4) and downstream at Fvb6_37.858Mb (LOD = 126.8), thus defining the limits of the 467kb female-function region (Fig 3). A QTL analysis of the seven linkage groups in homeologous group VI for which we had segregating markers in SDR vicinity amplicons (VI-B2-m not observed) supported the single QTL on ZW. The correlation between SDR genotype and female function was moderately high (R^2^ = 0.76; *P* < 10^−15^).

In EUR13×EUR3, we genotyped SDR-vicinity amplicons of 10 male-fertile and 10 male-sterile offspring. We observed 4 ZW segregating sites and 3 ZZ segregating sites in EUR13×EUR3 (Table S5; Fig S1). Similarly, PTR14×PTR19 we genotyped SDR-vicinity amplicons of 6 male-fertile offspring, 11 male-sterile offspring, and 3 offspring of unknown sex phenotype. We observed 12 ZW segregating sites and 4 ZZ segregating sites in PTR14×PTR19 (Table S5; Fig S1). Among both EUR13×EUR3 and PTR14×PTR19, all ZW SNPs in the SDR vicinity showed a perfect match to sex, with no observed recombinants.

### Recombination

Among all target capture linkage groups we observed 747 maternal recombination events and 731 paternal recombination events, suggesting a genome-wide recombination rate of 2.5 cM Mb^−1^ given a 700 Mb genome (Hirakawa *et al*., 2014). Recombination rates for 138 segments of 5-10Mb varied from 0 to 7.0 cM Mb^−1^ (median = 2.3, mean = 2.5, standard deviation = 1.4; Fig 4). There was no significant difference between maternal and paternal recombination rates genome-wide (Wilcoxon rank sum test, *P* > 0.1; Fig 4). Among the 56 linkage groups (28 from each octoploid parent), the entire ZW chromosome pair had the second-highest recombination rate estimate, 3.3 cM Mb^−1^ (Fig 4 pink circle), consistently higher than the rate in all other VI linkage groups (Fig 5), and only slightly exceeded by the I-Av-p rate (3.4 cM Mb^−1^, Table S8). Based on Fvb chromosome lengths, the ZW is estimated to constitute 2.3% of the *F. chiloensis* genome; the expected number of recombination events is therefore 34, assuming a constant rate and Fvb-proportional chromosome sizes, significantly lower than the observed 53 events observed in this chromosome (53:1425 ZW:non-ZW vs. 34:1444; χ2 = 10.9, P < 0.01). Conversely, the Z had one of the lowest recombination rates in males (1.7 cM Mb^−1^; Fig 4 blue circle), lower than all other VI linkage groups (Fig 5); only 5 linkage groups had lower rates (Table S8). Thus, the recombination rate for the sex chromosomes was nearly twice as high in females (ZW in the SDR) as in males (ZZ). (53:27 vs. 40:40 recombination events, χ^2^ = 8.5, *P* < 0.01).

**Fig 4.**
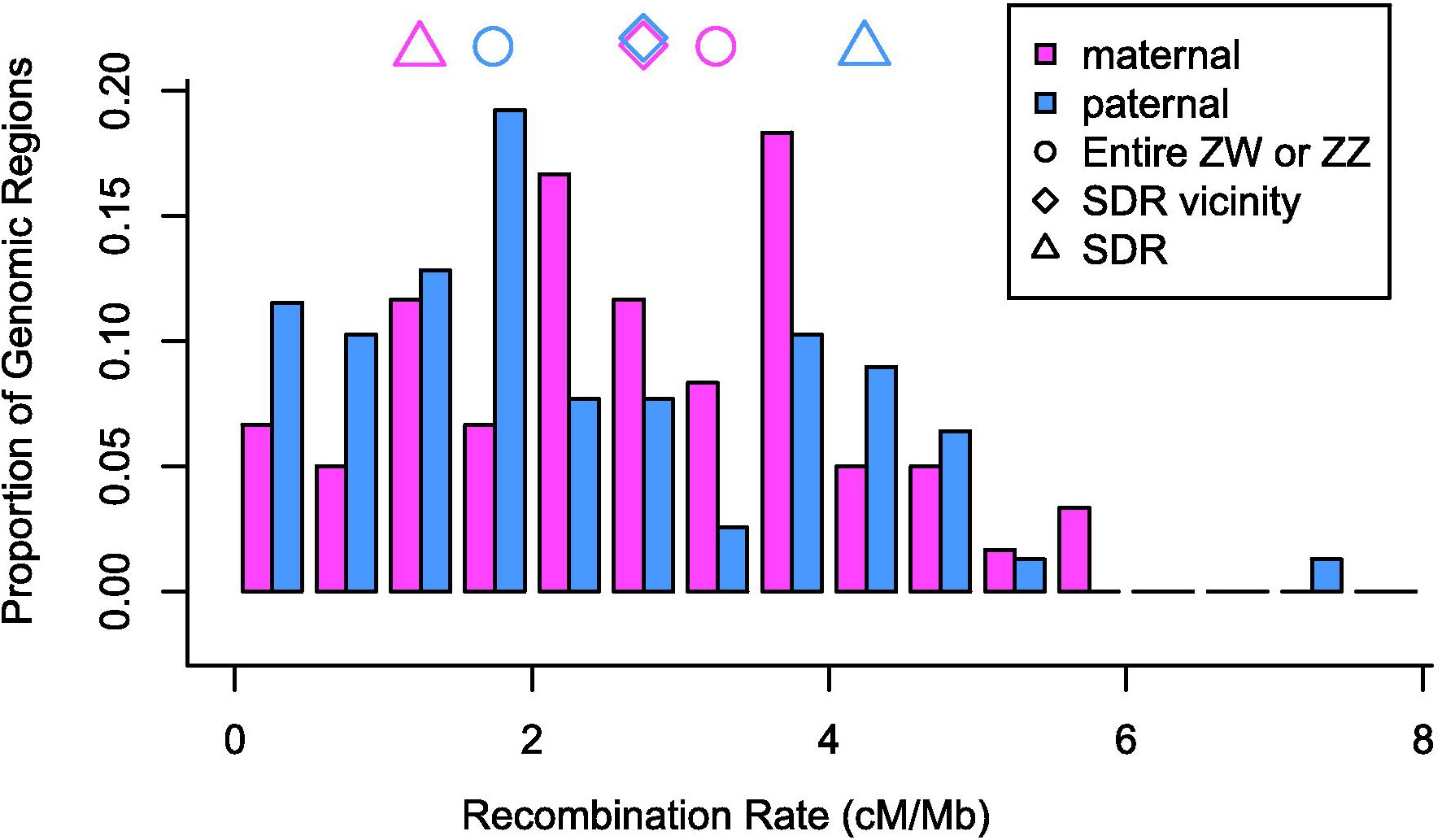
Genome-wide recombination rates in *Fragaria chiloensis* compared to ZW rates. The target capture linkage maps were divided into sections of 5-10Mb, free from any known rearrangements. Recombination rate was calculated for each section, as shown in this histogram of maternal (pink) and paternal (light blue) recombination rates. Symbols indicate maternal and paternal recombination rates across the entire sex chromosome (circles; ZZ or ZW), across the SDR vicinity from Fvb6_37.38–38.29 (diamonds), and across the SDR from Fvb6_37.38–37.71 Mb (triangles). Recombination is notably higher in the mother than in the father across the entire chromosome, but this pattern reverses at the SDR.

**Fig 5.**
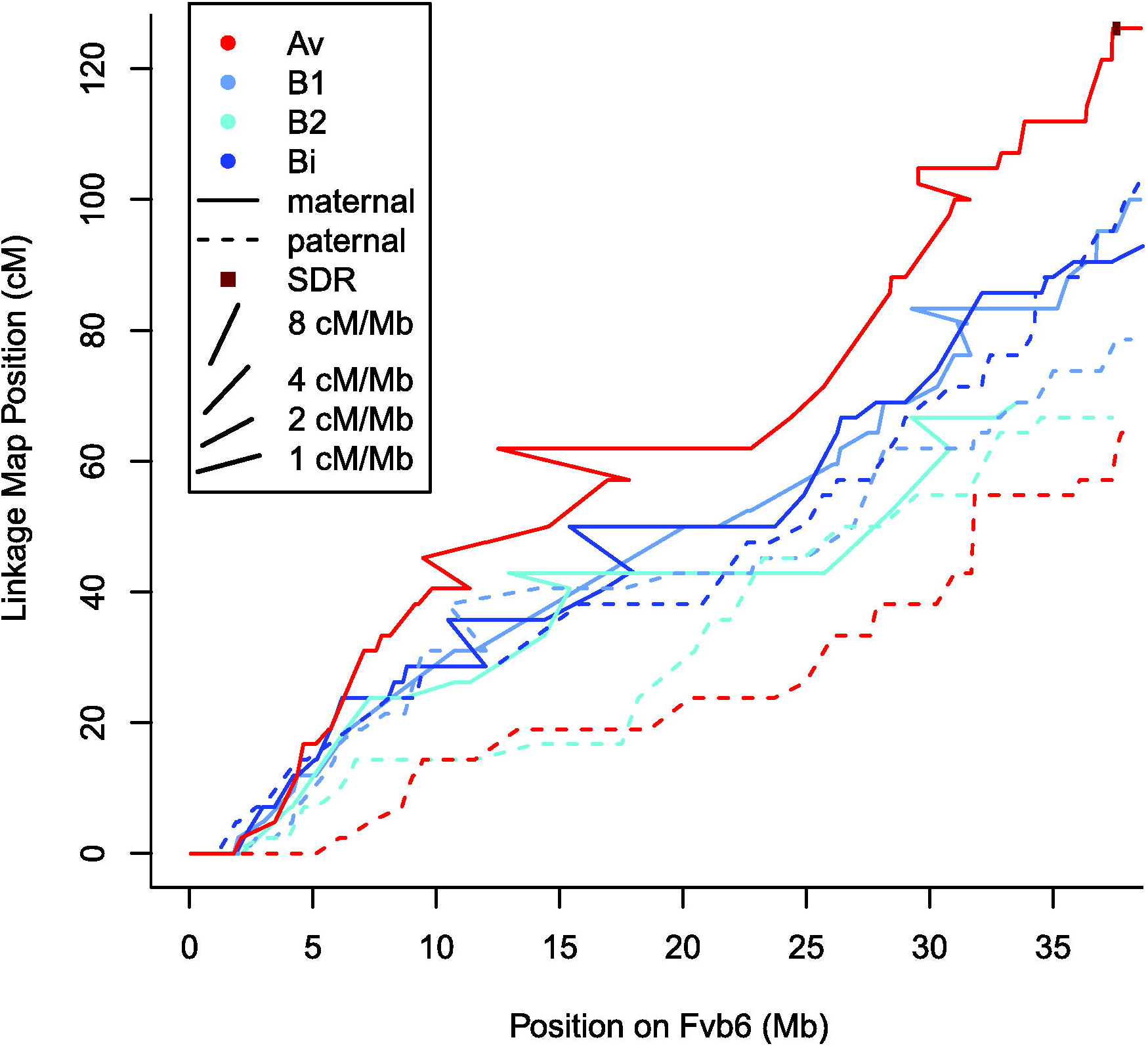
Recombination events across all eight *Fragaria chiloensis* VI linkage groups. Markers in each linkage group are plotted with physical Fvb reference genome position on the x-axis and centimorgan position on the y-axis. Recombination rates are therefore indicated by the slopes of the lines, as illustrated by the example slopes in the figure legend. Lines are color-coded based on subgenome, and are solid for maternal linkage groups and dashed for paternal linkage groups. The SDR is shown as a dark red box. For ease of visualization, markers showing radical rearrangements relative to the reference genome (<5% of all markers) have been removed. The ZW linkage group VI-Av-m harboring the sex determining region (SDR) has the highest recombination rate across the entire chromosome, while the ZZ linkage group VI-Av-p has the lowest recombination rate.

Across the 1Mb of the SDR vicinity in which we observed segregating sites in our amplicons, recombination rates were nearly identical in the maternal and paternal plants (2.9 cM Mb^−1^ and 2.8 cM Mb^−1^ respectively; Fig 4 diamonds). However, in the 330kb between Fvb6_37.378-37.708Mb, containing the entire male-function region and most of the female-function region, recombination was lower in the maternal parent (Fig 4 triangles), with six recombination events, implying a rate of 1.5 cM Mb^−1^, versus sixteen events (or 4.1 cM Mb^−1^) in the ZZ paternal parent (6:1209 vs. 16:1180 recombination events; χ^2^ = 4.7, *P* < 0.05). The recombination rate at the SDR in ZW plants was lower than the genome-wide average of 2.5 cM Mb^−1^, but not significantly (10 recombination events would be expected in the interval between Fvb6_37.378-37.708Mb, versus 6 observed; χ^2^ = 1.6, *P* > 0.1; Fig 4). Likewise, the recombination rate at the SDR in ZZ was higher than the genome-wide average but not an outlier. Unfortunately, we did not observe SNPs at both boundaries of this interval in any other VI homeolog, and therefore we cannot directly compare recombination rates among subgenomes specifically at the homeologous regions syntenic to the SDR.

### Divergence between W and Z

We observed no fixed differences between Z and W haplotypes, but many Z-W differences within crosses, suggestive of imperfect linkage disequilibrium with sex phenotype across the SDR vicinity. Specifically, in our three crosses, we observed 52 SNPs that differentiate the female parents’ Z and W haplotypes, most of them seen only in one family, not shared between the different crosses (HM1×SAL3: 36 SNPs; PTR14×PTR19: 12 SNPs; EUR13×EUR3: 4 SNPs; total distinct SNPs = 46; Table 2). This is a higher SNP density than for the same region compared between pairs of Z chromosomes (31 segregating in the father, not correcting for shared SNPs among crosses; Table 2; Fig S1). The highest density of W-specific SNPs occurred within the SDR, specifically in a 143kb “high W divergence” window between 37.565-37.708Mb, with 10 SNPs in coupling with male sterility/female fertility in HM1×SAL3, 4 in PTR14×PTR19, and 1 in EUR13×EUR3 (Table 2). For HM1×SAL3, the W-specific divergence in this high-divergence window was 0.39%, substantially higher than either the 0.07% of sites segregating on a single Z haplotype in this window (observed numbers of W versus Z SNPs 10:5, expected based on equal probabilities 3.75:11.25, χ^2^ = 13.9, *P* < 0.01), or the 0.06% W-specific divergence observed across the rest of the SDR vicinity. Thus, the divergence between the W and Z sequences within a cross appears particularly high in this window, especially for W-specific SNPs. Only one G/C SNP (here designated the “sex-associated marker”), at position Fvb6_37594072 in the high divergence window, was in coupling with male sterility/female fertility (W allele = C; Z allele = G) in all three crosses (Fig 3; Fig S1).

### Association between genotype and sexual phenotype in unrelated plants

In order to identify SNPs in linkage disequilibrium with the causal sex gene(s), we examined genetic diversity at SDR vicinity amplicons in 22 unrelated plants, 10 male-sterile and 12 male-fertile (the six parents of the crosses plus 16 additional plants). The sex-associated marker was heterozygous (G/C) in 9 out of 10 male-sterile plants (ZW), and homozygous (G/G) in 11 of 12 male-fertile plants (ZZ). Four additional SNPs (Fvb6_37565581, Fvb6_37565641, Fvb6_37567599, and Fvb6_37567962; Fig S1), all within the high W divergence window of the SDR, showed a weaker association with sex, but all were heterozygous in 80-90% of male-sterile plants and homozygous in 60-70% of male-fertile plants. Thus, of all observed SNPs, the single sex-associated marker showed the tightest linkage disequilibrium with the causal W haplotype.

## Discussion

As *F. chiloensis* possesses the youngest and least differentiated plant ZW sex chromosomes yet described, it illuminates how recombination rates may first begin to adjust in response to sex linkage. By extensively analyzing recombination, this study reveals rate heterogeneity patterns both expected and surprising, which could nonetheless be typical among the numerous dioecious plant species with homomorphic sex chromosomes. Our major results on *F. chiloensis* sex chromosomes indicate: (1) the SDR is physically small; (2) maternal PAR recombination is unusually high; (3) the SDR is characterized by SNPs in imperfect linkage disequilibrium.

### A small SDR

In contrast to most older sex chromosomes, the *F. chiloensis* SDR is smaller than 1 Mb, and recombination suppression is minimal. Most of the SDR vicinity has recombination rates typical of the rest of the genome, and nearly identical between ZW and ZZ plants (Fig 3). Both male and female function mapped to a single 280kb chromosomal region within the region first identified by Goldberg *et al*. (2010) at the distal end of the chromosome carrying the SDR. The estimated recombination rate in a 330kb SDR-centered window in a ZW parent is less than half of that in a ZZ parent, but is not unusually low relative to the rate for autosomes. Other unlinked loci or environmental effects may also influence female function, a quantitative trait for which the SDR explains 74% of variation, and which in other *Fragaria* can be highly environmentally labile (Spigler & Ashman, 2011) and influenced by additional minor-effect loci (Spigler *et al*., 2011; Govindarajulu *et al*., 2013; Ashman *et al*., 2012). Male function, in contrast, was almost perfectly determined by the SDR. Fewer than 1% of HM1×SAL3 offspring showed a mismatch, suggesting that any other factors affecting the propensity to produce pollen are negligible.

Male and female function may be controlled by two closely linked genes, or by the same gene. Presumably a single ZW gene initially controlled a single phenotype, likely male function given that gynodioecy, but not androdioecy, occurs in *Fragaria* (Li *et al*., 2012). Under the classic model of sex chromosome evolution (Charlesworth & Charlesworth, 1978), a second linked mutation occurs, yielding two distinct linked genes controlling male and female function. Recombination between these genes produces neuters, prompting strong selection for recombination suppression. In *F. chiloensis*, the two-locus model would require a remarkably small physical distance between the two loci (within the 280kb SDR), but this could arise either coincidentally or via adaptive translocation of the second locus in order to achieve tight linkage. Because recombination at autosomal rates should only occur rarely within 280kb (fewer than 1% recombinant offspring given 2.5 cM Mb^−1^ genomic mean rate), selection for additional recombination suppression would not be particularly strong. Although recombination between loci controlling male and female function would theoretically produce neuters (Charlesworth, 2015), we observe only 7 out of 619 HM1×SAL3 offspring showing both male and female sterility, none of which are recombinant on ZW, and four of which produced only 1-2 flowers and may have failed to fruit by chance. Alternatively, there might be only a single sex determinant at the SDR, as previously hypothesized for *Fragaria* (Ahmadi & Bringhurst, 1991) and other dioecious systems with only a single identified sex regulator (Akagi *et al*., 2014). Variation in female function among male-fertile individuals in a gynodioecious ancestor could have been influenced by gene(s) anywhere in the genome, not necessarily on ZW/ZZ. If males had higher fitness than hermaphrodites (Charlesworth, 1989), alleles conferring female sterility (but only in male-fertile individuals due to sex-specific gene expression; Pennell & Morrow, 2013) would increase in frequency or even fix, leading to the highly dioecious condition observed in *F. chiloensis* even without new sterility mutations on ZW/ZZ. Selection could favor linkage of female fertility modifiers to the SDR, but such selection would be weak if these mutations had little phenotypic effect in females (Pennell & Morrow, 2013). If a single gene controls both male and female function in *F. chiloensis*, there would only be selection for recombination suppression if other linked genes experienced sexually antagonistic selection (reviewed in Otto *et al*., 2011). Alternatively, reduced recombination could be a neutral consequence of high sequence divergence (Opperman *et al*., 2004).

Several promising candidate causal sex gene(s) occur among the 70 known SDR genes (Table S7). Genes in families previously implicated in sex function include a mitochondrial-targeted pentatricopeptide repeat protein (Chen & Liu, 2014), two methyltransferases (Geraldes *et al*., 2015), and a cluster of cystinosin homologs (Besouw *et al*., 2010). An important future direction will be to test SDR genes for sexual function or sexually antagonistic alleles.

### Recombination at the PAR

In addition to suppressed recombination at the SDR, sex chromosomes frequently show elevated recombination in the PAR, and thus typically a chromosome-wide average rate similar to autosomes (reviewed in Otto *et al*., 2011). In *F. chiloensis*, the PAR represents over 99% of the 39Mb sex chromosome. Thus, the PAR recombination rate is approximately equal to the chromosome-wide average rate, which remarkably is nearly twice as high for ZW than for ZZ.

Recombination rates are lower on the other *F. chiloensis* VI linkage groups, and are lower among all 56 *F. virginiana* linkage groups (Tennessen *et al*., 2014), suggesting that the high ZW rate in *F. chiloensis* is newly evolved. Although elevated PAR recombination has been attributed to the need for least one crossover within small PARs (Otto *et al*., 2011), this cannot explain high rates in large PARs. Thus, the adaptive value of the elevated *F. chiloensis* recombination rate, if any, is not to ensure a minimum chromosome-wide rate. Instead, perhaps there is a benefit for certain alleles to recombine away from the SDR, at least in the early stages of sex chromosome evolution. Plastic recombination can be adaptive if recombination modifiers can detect *cis-trans* effects (Agrawal *et al*., 2005), although testing the feasibility of this scenario in *F. chiloensis* specifically would require extensive modeling. Because the W may accumulate deleterious mutations due to low effective population size and/or exhibit lower fitness per se than the Z, as suggested by the fluctuating, male-biased sex ratios observed in *F. chiloensis* and other dioecious plants (Hancock & Bringhurst, 1980; Field *et al*., 2013), elevated recombination could hypothetically break up deleterious linkage disequilibrium between the PAR and the SDR.

Examination of recombination rate is standard in studies of sex chromosomes (Charlesworth, 2015), but the statistical detection of recombination suppression or elevation is not always straightforward. Although integrating linkage maps with reference sequences is increasingly common (Wang *et al*., 2012; Zhang *et al*., 2015; Papadopulos *et al*., 2015) the resultant recombination comparisons still carry several caveats. First, recombination rate varies across the genome, so inferring suppressed or elevated recombination depends on what control is used (Nachman, 2002; Natri *et al*., 2013). For example, although maternal recombination is lower at the SDR than the PAR (Fig 4), this is due at least as much to elevated recombination at the PAR (relative to the autosomes) as to suppressed recombination at the SDR. Second, major rearrangements such as large insertions or deletions relative to the reference genome (in this case from *F. vesca)* would affect recombination estimates. The 0.7 Gb octoploid *Fragaria* genome has likely undergone some gene loss and other rearrangements relative to the 0.2 Gb diploid reference (Hirakawa *et al*., 2014). However, we minimized the effects of this limitation by identifying and avoiding rearrangements in the target capture map, by focusing on the subgenome (Av) most closely related to *F. vesca*, and by making direct comparisons between maternal and paternal maps under the assumption that most large rearrangements would be present in both parents.

### Linkage disequilibrium at the SDR

The genetic basis of sex determination is highly variable among *Fragaria* species, often mapping to different locations on chromosome VI (Goldberg et al. 2010; Spigler et al., 2010, 2011; Ashman et al. 2015) or other chromosomes (Tennessen et al. 2013). In contrast to this interspecies variability, the SDR is highly consistent within *F. chiloensis*. The same SDR was identified in three *F. chiloensis* crosses from populations up to 1000 km away, suggesting it may be the most common, or only, SDR in this species, at least in North America. A single “sex-associated marker” was in coupling with male sterility/female fertility phenotype in all three crosses, and this SNP also showed near-perfect association with sex across a set of unrelated plants. Of the two exceptions, one plant was the sole specimen from South America but the other was from the same population as one of the crosses (PTR14×PTR19), suggesting either that the SDR does not perfectly control sex in all cases, or that the sex-associated marker is not in perfect linkage disequilibrium with the causal gene(s). One consequence of recombination suppression should be that a W-specific haplotype behaves as a single unit in population genetics, with multiple SNPs in linkage disequilibrium with sex. The fact that only a single sex-associated marker was observed, despite our sequencing 2% of the 280kb SDR, suggests that there are few SNPs (<100) showing such high linkage disequilibrium with sex. In turn, this observation suggests that the true SDR may be substantially smaller than the 280kb mapped region.

We observe a notably high concentration of maternally-heterozygous SNPs in a small genomic section corresponding to the second half of the SDR (Fig 3, Fig S1). Other than the sex-associated marker, these SNPs are not W-specific across populations, but may show moderate linkage disequilibrium with the W haplotype. Thus, the SDR is characterized by variants showing imperfect associations with sex owing to several potential reasons: suppressed recombination, selection against recombinants, or low effective population size at the SDR due to fluctuating sex ratios, such that deleterious mutations are not purged by selection (Ohta 1973; Wang *et al*., 2012; Geraldes *et al*., 2015). High PAR recombination may in fact have evolved to prevent maladaptive linkage to W-associated mutations (Agrawal *et al*., 2005). These processes could lead to even greater Z-W differentiation. Alternatively, given that non-recombining sex chromosomes seem to have only rarely become established in dioecious flowering plants, they may not evolve in *F. chiloensis*, and Z-W divergence could remain low indefinitely, especially if recombination eliminates W-specific changes.

## Acknowledgements

We thank C. Ashman, T. Ashman, R. Dalton, C. Kustek, K. Mazzaferro, B. Rosa-Neves, K. Schuller, L. Stanley, H. Wipf, E. York, and the University of Idaho IBEST Core Facilities staff for glasshouse, field, laboratory, or data assistance, and the Ashman and Liston lab members and three anonymous reviewers for helpful comments. This work was supported by the University of Pittsburgh and the National Science Foundation (DEB 1020523, DEB 1241006, and RET/REU supplements to T-LA; DEB 1020271 and DEB 1241217 to AL).

## Author Contributions

Design of the research: T-LA, AL; Performance of the research: RG, T-LA; Data analysis, collection, and interpretation: JAT; Writing the manuscript: JAT, T-LA, AL, and RG.

## Supporting information captions

**Table S1.** IDs and collection localities of plants unrelated to the parents of the crosses.

**Table S2.** Primers for amplicon sequencing.

**Table S3.** PCR conditions for all amplicons.

**Table S4.** Phenotypes and genotypes for all HM1×SAL3 offspring.

**Table S5.** Phenotypes and genotypes of EUR13×EUR3 and PTR14×PTR19 offspring.

**Table S6.** Positions of all markers in target capture linkage maps.

**Table S7.** List of 70 genes in the 280kb window on chromosome Fvb6 matching the *F. chiloensis* SDR.

**Table S8.** Recombination rates for all linkage groups.

Fig S1. Haplotypes in the sex determining region (SDR) vicinity of *Fragaria chiloensis*.

